# In-Depth Immune Profiling Reveals Cell-Type, Sex and Infection-Stage Dependent Differences for Tuberculosis Control

**DOI:** 10.64898/2026.09.30.755733

**Authors:** Jacqueline Watt, Dustin J. Sokolowski, Jun Liu

## Abstract

Tuberculosis (TB) disease consistently causes the highest number of annual deaths from infectious disease and is responsible for over 100 million disability-adjusted-life-years annually. Despite the dramatic impact on human health, TB treatments are intensive and moderately effective, while Bacille Calmette-Guérin (BCG), the primary TB vaccine strategy and the oldest vaccine in circulation, needs updating. One gap in knowledge is that the vaccine primed lung immune environment has not been systematically profiled following TB challenge, making it challenging to interpret how various vaccine strategies provide protection. Here we profiled the male and female mouse lung immune environment using Mass Cytometry by Time of Flight (CyTOF) and bacterial burden (colony forming units, CFU), to systematically identify immune cells contributing to *Mycobacterium tuberculosis* (*Mtb*) infection or vaccine-induced protection. We investigated the mouse immune system after various infection outcomes including primary *Mtb* infection, *Mtb* reinfection, *Mtb* challenge after BCG vaccination and *Mtb* challenge after vaccination with a live attenuated *Mtb* vaccine. Across all conditions we identified Thy1.2 expression on NK cells as a new candidate for a sex-specific correlate of protection. Our results further replicate and support known immune responses for vaccine-induced protection, such as T-cell activation and infiltration. These findings highlight the pan-immune response to vaccine-induced protection, showcase the importance of sex in rodent models of TB, and identify novel host immune candidates to protect the lung against TB disease.

## Introduction

Tuberculosis (TB), caused by *Mycobacterium tuberculosis* (*Mtb*), is an infectious disease responsible for over one million deaths and ten million new cases annually, with roughly 1.7 billion individuals harboring latent TB infection (1). Outside of the COVID-19 pandemic, TB has consistently caused the highest number of deaths from any infectious disease for decades. Beyond mortality, TB imposes a substantial burden on quality of life, accounting for an estimated 122 million disability-adjusted life-years from fatal and non-fatal health loss in 2019 alone (2).

Treatment of acute TB typically involves a two-month “intensive phase” of high-dose combination therapy with Isoniazid, Rifampicin, Pyrazinamide, and Ethambutol, followed by a four-month “continuation phase” of Isoniazid and Rifampicin alone (2HRZE/4HR). These regimens are associated with significant adverse effects; in some populations, nearly 20% of individuals experience serious drug reactions, and approximately 2% are unable to complete therapy due to these effects (3). Moreover, antibiotic resistant strains of *Mtb* continue to emerge, underscoring the critical importance of vaccination as a preventative strategy against TB (4).

The Bacille Calmette-Guérin (BCG) vaccine, derived from *Mycobacterium bovis*, is currently the primary TB vaccination strategy (5). First administered to humans in 1921, BCG is the oldest vaccine still in use and remains the most widely administered worldwide, with roughly half of the global population vaccinated (6). Despite its widespread use, BCG efficacy varies widely across populations (6–10). Protection is generally higher in individuals aged 0–15, but multiple boosting strategies have struggled to improve efficacy in adults (8,11). Sex also influences TB susceptibility; males exhibit higher infection risk and immunoreactivity than females, a disparity that increases with age (12). Sex differences in TB severity are observed in mice as well (13), with recent work attributing these differences to variations in the interplay of the adaptive and innate immune responses in each sex (14).

*Mtb* has infected humans for an estimated 75,000 years (15), and new strains with differing fitness continue to arise (16). This long-term evolutionary arms race has led to intricate interactions between *Mtb* and the human immune system, engaging multiple cell types in the airway epithelium, innate immunity, and adaptive immunity (17). Consequently, live attenuated vaccines derived from *Mtb* may provide improved protection by stimulating immune responses more analogous to natural infection. One promising candidate, MTBVAC, carries deletions in the virulence genes *phoP* and *fadD26* (18). MTBVAC has demonstrated safety and protective efficacy in animal models and is currently undergoing Phase 3 clinical trials (19).

A better understanding of how *Mtb*, BCG, and attenuated vaccines interact with the immune system is essential for improving TB vaccination strategies. There continues to be a large effort to identify and validate correlates of protection, recently of note is the TB immune correlates program established for BCG and another TB vaccine currently in Phase 3 clinical trials, M72/AS01_E_, induced correlates of protection (20). We have previously described an attenuated *Mtb* strain by knocking out the *lsr2* gene (Δ*lsr2*) (21), which functions as a global transcriptional repressor of multiple virulence associated genes (22–25). In that same study we also showed mice inoculated with Δ*lsr2* displayed an intermediary immune activation and exhaustion pattern between those observed with the parental *Mtb* strain and BCG using Mass Cytometry by Time of Flight (CyTOF) in female mice (21). Of the 10 lung immune cell types observed, all of them exhibited changes in either proportion or expression of activation and/or exhaustion markers, reflecting a broad and highly dynamic immune response to mycobacterial infections (21). Matched immune profiling of lung tissue from primary infected, vaccinated, and reinfected animals will provide a more complete picture for the immune response to *Mtb*.

CD4 and CD8 T cell infiltration and activation correlate with vaccine-induced protection, although multiple other cell types also contribute (26). Despite significant progress, many vaccine-induced correlates of protection remain undefined (26). A systematic, high-dimensional screening of lung immune responses allows direct comparison of immune effect sizes within a single controlled experimental environment. CyTOF enables single-cell resolution profiling of up to twenty samples simultaneously, facilitating identification and comparison of correlates of protection across vaccine strategies and immune cell types in *Mtb* challenge studies.

Here we analyzed the immune profiles of male and female mice after *Mtb* challenge following several vaccination strategies. Our results recapitulate well established findings, such as T cell activation serving as a potential correlate of protection (27). Additionally, we identified novel trends, such as Thy1.2 expression on NK cells being a strong correlate of protection in female mice despite having no correlation in males. Collectively, these findings expand our understanding of how vaccination and *Mtb* infection shape the lung immune landscape and reinforce the need to include male and female sexes in future work to address the unique responses of each sex and improve our overall approach to TB treatment and prevention.

## Results

### Female mice display better long-term protection with both vaccine-induced and natural immunity to Mtb infection

We first aimed to profile the immune system in the lung at various stages pre and post vaccination. We used CyTOF to systematically profile lung immune cells in the mouse at a single-cell resolution. Groups of male and female C57BL/6 mice were inoculated with either Δ*lsr2*, BCG-Japan or PBS as a sham control. We previously described Δ*lsr2* as an effective vaccine strategy for *Mtb* infection (28). BCG-Japan was selected as it has been shown to be more immunogenic and more effective at reducing risk of TB in infants compared to other BCG strains (29–31). An additional group was also infected with *Mtb* H37Rv for 4 weeks and then treated with rifampicin and isoniazid for an additional 12 weeks (*Mtb*+antibiotics[abx]) following a previously established protocol (32). Thus, Δ*lsr2* and BCG represent vaccine-induced immunity conditions, and the *Mtb*+abx group represents a natural immunity condition. 18 weeks after the initial inoculations, all groups were aerosol challenged with ∼100 colony forming units (CFUs) of *Mtb* H37Rv using the Glass-Col Inhalation Exposure System (Figure 1A). We profiled three infection stages: ‘baseline’, 18 weeks post-vaccination and pre-challenge with *Mtb*; ‘early infection’, 3 weeks post-challenge with *Mtb;* and ‘late infection’, 12 weeks post-challenge with *Mtb*. We measured bacterial burden (CFU) in the lungs and spleen of each mouse as well as collected single cell suspension samples for immune analysis.

**Figure 1.**
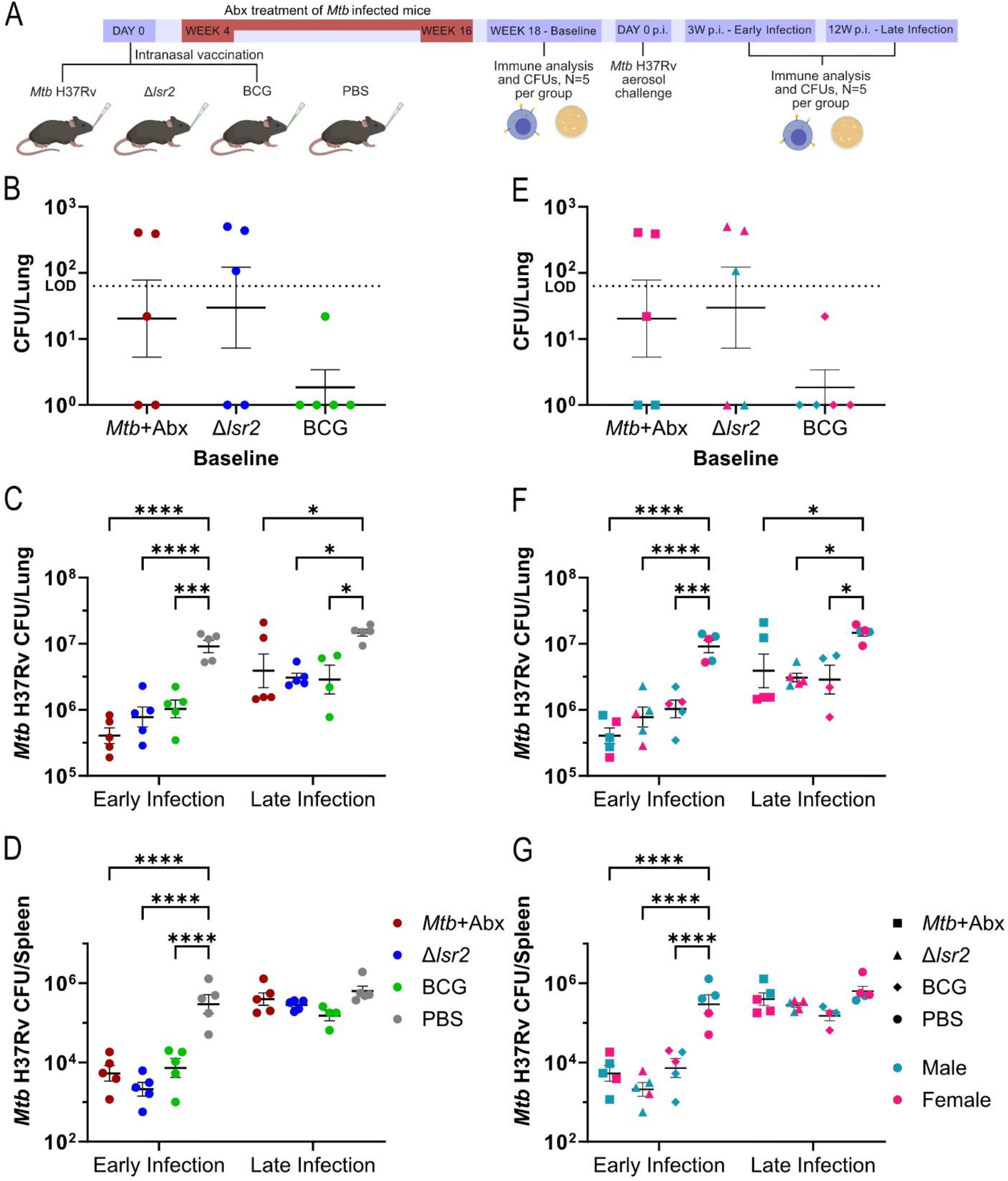
Long term protection to *Mtb* by multiple vaccination strategies in male and female C57BL/6 mice. (A) Schematic representation of experimental design. 5 C57BL/6 mice per group, with 2-3 males and females per group, were intranasally vaccinated with 5×10^5^ CFU of Δ*lsr2* or BCG in 20μL PBS, or 5×10^3^ CFU of *Mtb* H37Rv. After 4 weeks, mice inoculated with *Mtb* H37Rv were treated with 200mg/L of rifampicin and 100mg/L of isoniazid in the drinking water for 12 weeks. ‘Baseline’ CFUs of the bacterial strains used for vaccination in the (B) lungs were determined pre-challenge with *Mtb*. Limit of Detection (LOD) = 65. 18 weeks after vaccination, all mice were aerosol challenged with *Mtb* H37Rv (∼100 CFU/lung). All groups were followed for up to 12 weeks and *Mtb* H37Rv CFUs in the (C) lungs and (D) spleen was determined at ‘early infection’, 3 weeks post-challenge, and ‘late infection’, 12 weeks post-challenge. (E-G) shows the bacterial burden data overlayed with the mice sexes. Male mice are teal, and female mice are pink. Data presented as mean ±SEM. Statistical significance was determined by two-way ANOVA with Tukey’s multiple comparisons test (*, p<0.05; **, p<0.01; ***, p<0.001; ****, p<0.0001; ns = not significant).

At ‘baseline’, as expected, we found a very low number of CFUs in the lungs and there were no detectable bacteria in the spleen for the BCG, Δ*lsr2* and *Mtb*+abx groups. In the lungs, six out of 10 of the *Mtb*+abx and Δ*lsr2* vaccinated mice and 1 out of 5 BCG vaccinated mice had some bacterial burden after 18 weeks (Figure 1B). At the ‘early infection’ timepoint, the PBS control groups had a significantly higher bacterial burden compared to all vaccinated groups in both the lung and spleen (Figure 1C-D). Bacterial burden was also highly correlated between the lung and spleen (R^2^ = 0.751, p = 1.33 x 10^-4^). We did not find sex differences for bacterial burden in the lungs or spleen at the ‘early infection’ timepoint (Figure 1F-G). At the ‘late infection’ timepoint PBS control mice had a greater bacterial burden than vaccinated mice in the lung; bacterial burden was also correlated between the lung and spleen (R^2^ = 0.743, p = 2.69 x 10^-4^; Figure 1C-D).

Unlike at ‘early infection’, we found considerable sex differences based on vaccination strategy at the ‘late infection’ timepoint (Figure 1F-G). In the lungs we found sex differences in bacterial burden with BCG (FDR = 0.013) and *Mtb*+abx (FDR = 2.55×10^-4^), where male mice had a higher bacterial burden than females. We did not observe sex differences in the Δ*lsr2* vaccinated group. Moreover, *Mtb*+abx male mice had a bacterial burden that more closely reflected PBS control mice, with a log_10_CFU > 7. Additionally, *Mtb*+abx male mice had a higher bacterial burden than Δ*lsr2* male mice (FDR = 0.018) and the high bacterial burden of *Mtb*+abx males also reflected a vaccination-by-sex interaction on bacterial burden (FDR = 0.048). Together, we observed a strong effect of vaccinated versus unvaccinated mice at the ‘early infection’ timepoint, and a sex- and vaccine-strategy-specific response for bacterial burden at the ‘late infection’ timepoint in both the lungs and spleen. The linear relationship between lung and spleen bacterial burden is well characterized during *Mtb* infection and is consistent across all groups and sexes (33). As such, we focused on lung bacterial burden as the primary outcome variable for CyTOF analyses, rather than spleen bacterial burden.

### Profiling of mouse lung immune cell abundances over 30 weeks using multiple TB vaccine strategies

CyTOF analysis with 28 immune cell proteins was performed and 10 major immune cell subsets were identified. Each timepoint was treated as a different experimental batch with 59 mice in total. After quality control and filtering, we collected 701,654 total cells (mean 36,929 cells per sample), 978,879 total cells (mean 48,944 cells per sample), and 1,469,319 total cells (mean 77,333 cells per sample) at the ‘baseline’, ‘early infection’, and ‘late infection’ timepoints respectively. We adopted a cell-type label transferring pipeline developed for single-cell RNA-seq to our dataset to computationally match cell-types between timepoints, as each timepoint represents a different CyTOF experiment. This approach allowed us to perform one-to-one mapping of cell types between timepoints, but not of clusters and metaclusters, which reflect cell subtypes and states (Supplementary Figure S1). As such, we focused on differential cell-type abundance between timepoints.

Broadly, cell-type abundances followed the expected patterns based on our previous immune profiling work in the female *Mtb* infected lung 6 weeks post infection (21). Like bacterial burden, cell-type abundance was the most dynamic as a function of timepoint, followed by vaccine strategy and sex. Briefly, when considering all samples together, every cell-type was differentially abundant across timepoints, reflecting the global immune reprogramming resulting from *Mtb* infection (Figure 2A). CyTOF let us compare relative and not absolute cell-type abundance, so the decrease in some cell-type proportions during ‘early infection’ may reflect a lower proportion of a cell-type that did not infiltrate the lung, rather than a loss of that cell-type in the lung. NK cells, other T cells, and Tregs were conditional on sex (Figure 2B). Granulocyte and possibly CD4 T cell abundance were biased by vaccine strategy (Figure 2B). CD4 T cell activity and infiltration are well characterized as being critical for protection against *Mtb* infection (27). Granulocytes, in particular neutrophils, on the other hand are understudied and their role in providing a protective immune response versus a permissive response is contested, despite being one of the first cells to arrive at the site of infection and being a main cell-type found in granulomas in humans (27).

**Figure 2.**
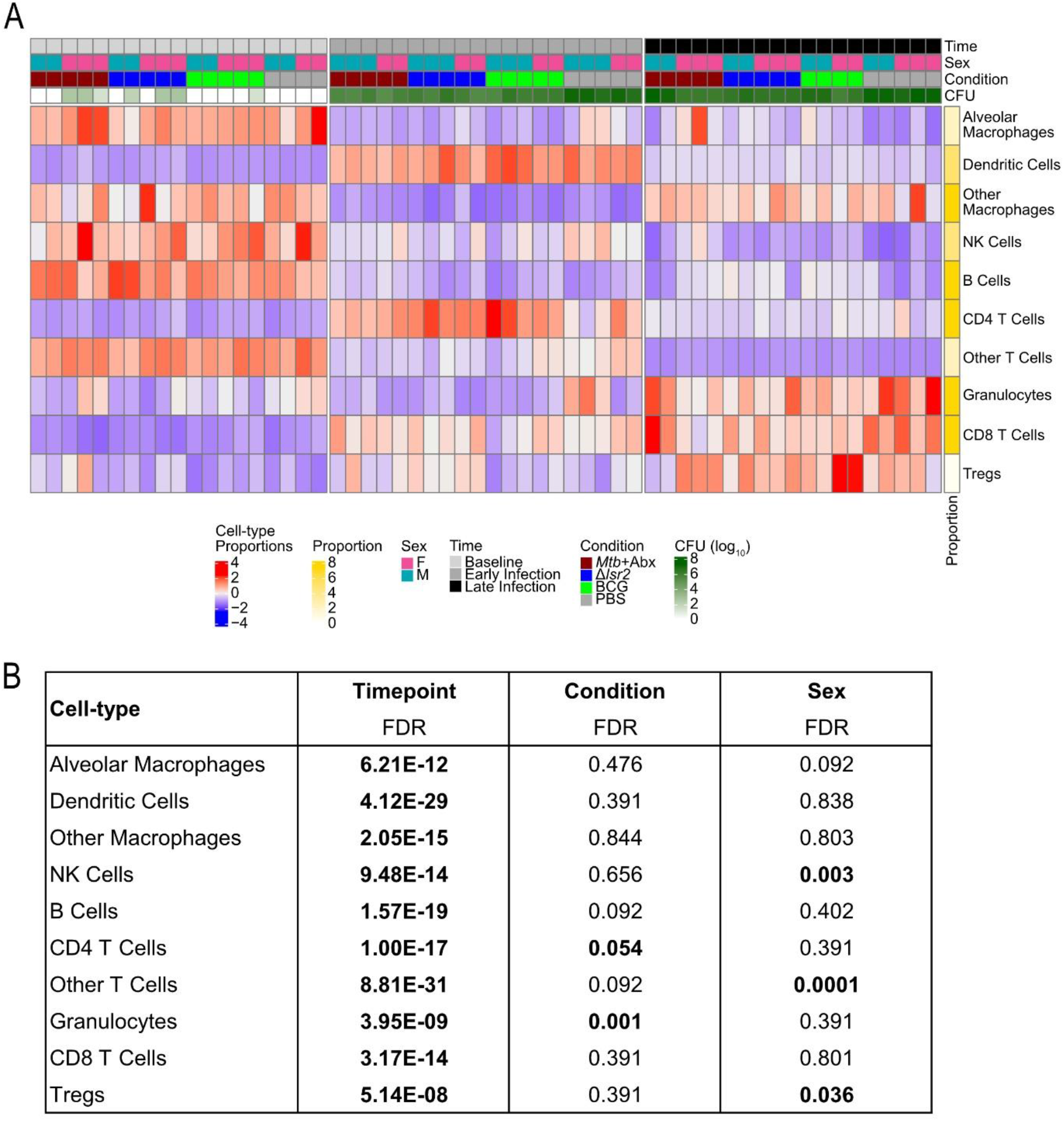
Global summary and cell-type composition of the male and female lung immune system after various vaccine strategies pre and post *Mtb* challenge. (A) Heatmap of row-normalized cell-type proportions for every sample included in each CyTOF experiment. Rows are cell-types and columns are individual samples. Column annotations represent timepoint, condition, sex, and bacterial burden log_10_CFU. Row annotations represent the median cell-type proportion across all samples. (B) Table of cell-type differential abundances that are associated with timepoint, condition or sex. Significant associations (FDR ≤ 0.05) are in bold.

Each measured timepoint represents a different stage of infection, as such, the variance introduced by timepoint makes it difficult to interrogate the variation across vaccine strategy and sex when looking at all timepoints simultaneously. We next investigated differential cell-type abundance at each timepoint individually (Figure 3A). As expected, we did not find differences in cell-type abundance at the ‘baseline’ timepoint. In contrast, we found that aerosol *Mtb* challenge led to large scale immune reprogramming of multiple cell types in a vaccine strategy and sex dependent manner during ‘early infection’ (Figure 3B). Here, we focused on cell types whose abundance was associated with bacterial burden (log_10_CFUs) and was differentially abundant across vaccine strategies. Specifically, other T Cell proportions were associated with bacterial burden and differentially abundant between vaccine strategy and sex. Granulocyte proportions were also associated with log_10_CFU, and differentially abundant between vaccine strategy, sex, and their interaction. Lastly, alveolar macrophage proportions were associated with log_10_CFU, and differentially abundant between vaccine strategy.

> ‘Late infection’ immune cell compositions paralleled ‘early infection’ in that they were also correlated with bacterial burden. Specifically, other T cells, Tregs, NK cells, B cells, dendritic cells, and alveolar macrophages were negatively associated with bacterial burden, and CD8 T cells and granulocytes were positively associated with bacterial burden during ‘late infection’ (Figure 3C). Other T cells, Tregs, and NK cell proportions were all also associated with sex and possibly associated with vaccine strategy.

**Figure 3.**
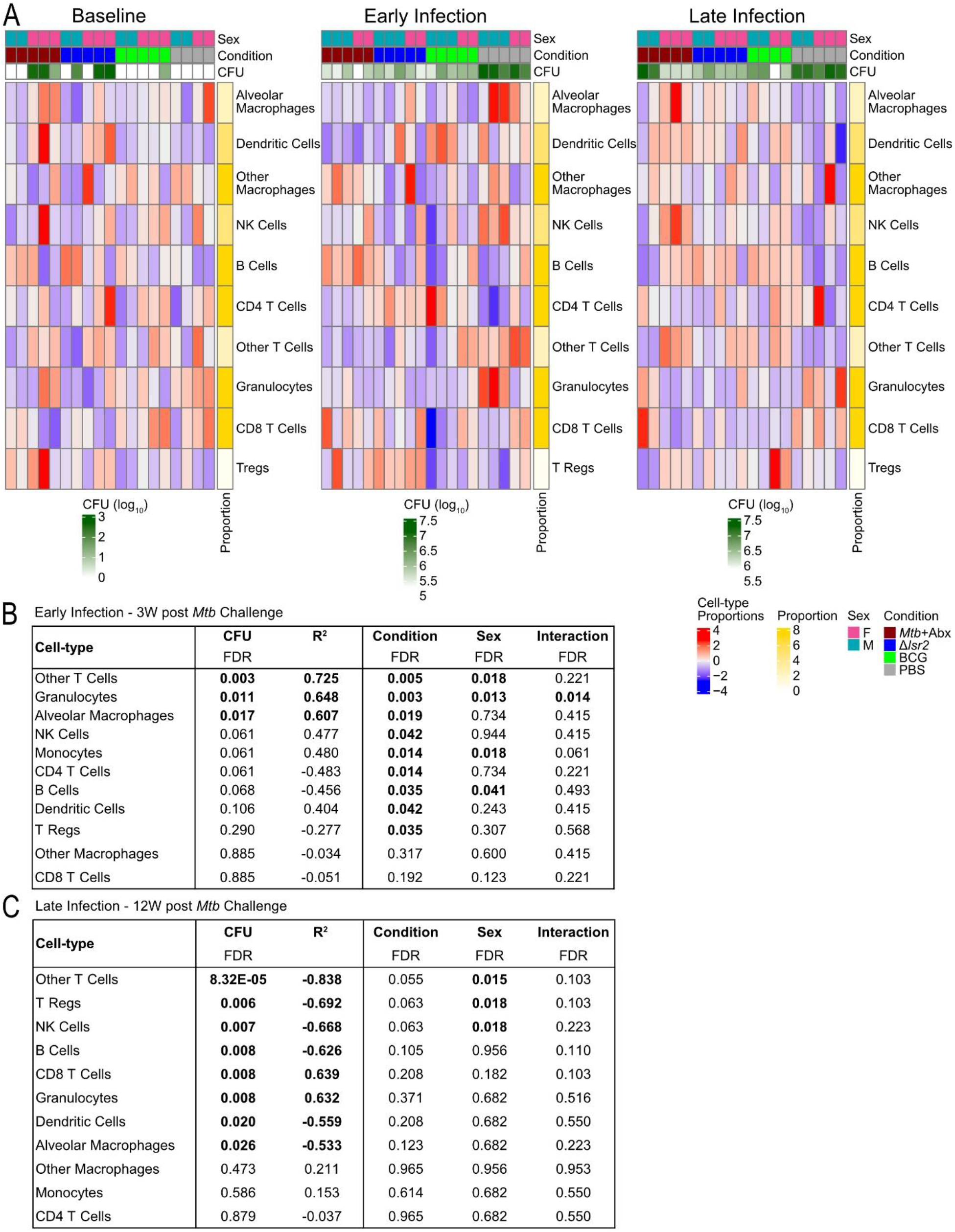
Cell-type differences within vaccine strategies and sexes. (A) Heatmap of cell-type proportions at each timepoint scaled by cell-type. Rows are cell-types and columns are individual samples. Column annotations represent timepoint, condition, sex, and bacterial burden log_10_CFU. Row annotations represent the median cell-type proportion across all samples. Table of cell-types that are associated with bacterial burden and are differentially abundant across condition or sex during (B) ‘early infection’ (3W post *Mtb* challenge) and (C) ‘late infection’ (12W post *Mtb* challenge). Significant associations (FDR ≤ 0.05) are in bold.

### Δlsr2 vaccinated and Mtb+abx female mice show differential adaptive and innate immune cell activation 18 weeks after inoculation

After computing differences in cell-type abundance within and between timepoints, we focused on cell-type-specific protein expression associated with bacterial burden, condition, and sex (Figure 4A) using an approach focusing on Principal Component Analysis (PCA). PCA is often used as a quality control metric in ‘omic’ datasets including CyTOF (35) because, for example, if plotting samples along PC1 separates them by condition, then the largest proportion of variance in the dataset is driven by the experimental design (36). When analyzing PCA, the datapoint that each mouse has on each principal component is called their PC score, and the weight that a cell-type-specific protein contributes to each principal component is called the PC loading (34). By analyzing PC scores, we can ask how the associated vaccine strategy and sex are related to the PC, and by analyzing PC loadings, we can ask which cell-type-specific proteins are driving the principal component.

**Figure 4.**
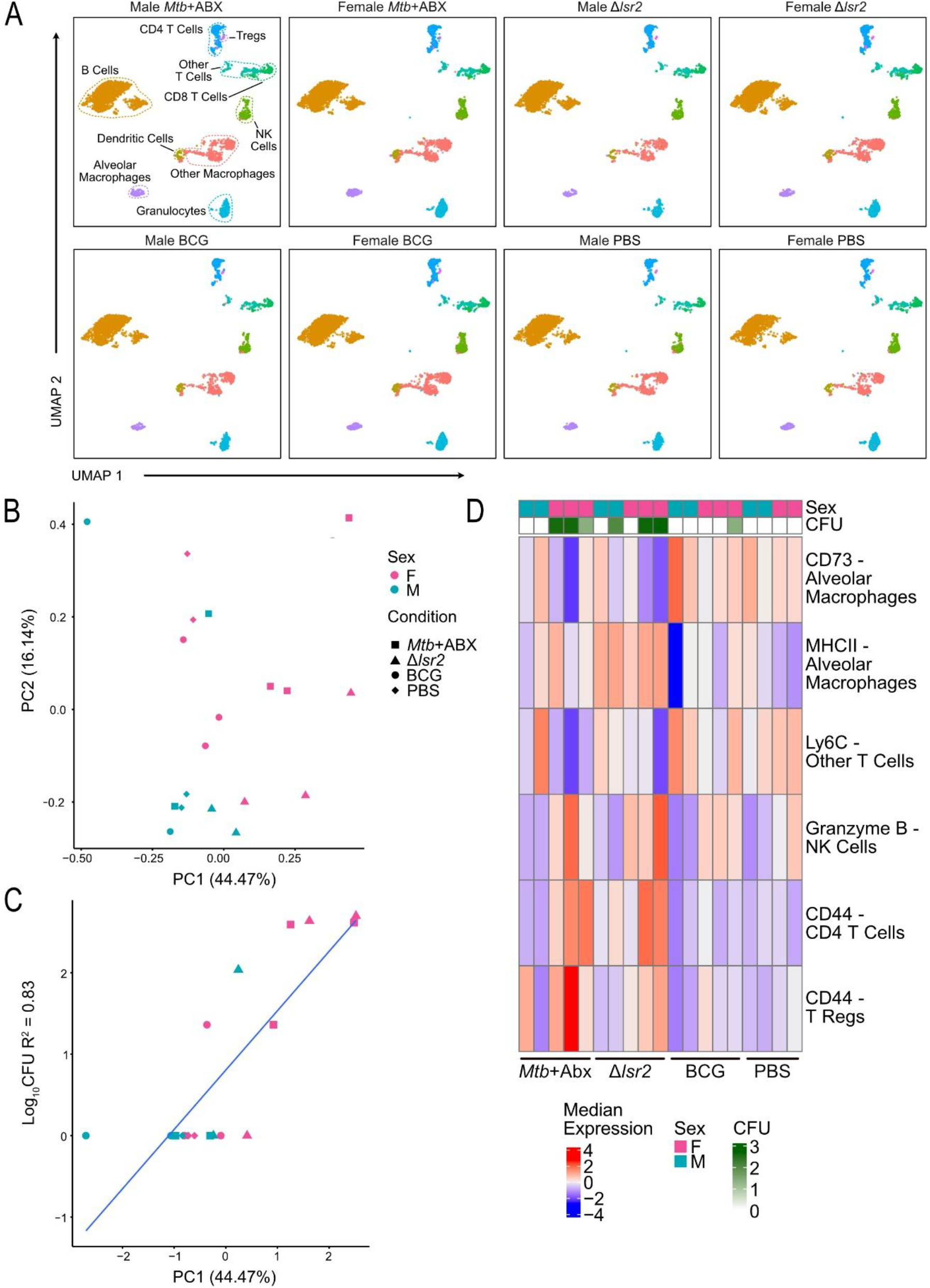
Immune profile of the male and female mouse lung 18 weeks after various mycobacterial inoculation strategies. (A) Uniform manifold approximation mapping (UMAP) showing the distribution of immune cells in the lungs based on their protein profiles, separated by treatment and sex. (B) Scatterplot of the first two principal components (PC) from a PC analysis of pseudobulked protein expression profiles for all samples. Shape represents condition and colour represents sex. (C) Scatterplot showing the relationship between PC1 scores (x-axis) and bacterial burden (log_10_CFU, y-axis). Conditions are shape and sex is colour. (D) Heatmap of cell-type-specific protein expression driving PC1. Rows are cell types, columns are samples, and heat is the median expression of pseudobulked protein expression in each cell-type.

At the ‘baseline’ timepoint, pseudobulked PCA showed that female mice in both Δ*lsr2* and *Mtb*+abx condition groups were linearly separable across PC1 (44.47% of variation), and there was no clear bias of vaccine strategy or sex across PC2 (16.14% of variation; Figure 4B). As such, we found that the background strain used for the initial inoculation (*Mtb* vs. *M. bovis*) influenced immune reprogramming in female mice.

PC1 was also associated with bacterial burden at ‘baseline’, as well as condition and sex (Figure 4C). We next sought to investigate cell-type-specific protein expression driving PC1. We found that CD73 in alveolar macrophages and Ly6C in other T cells being negatively associated with bacterial burden, and MHCII in alveolar macrophages, CD44 in CD4 T Cells, CD44 in T regs, and Granzyme B in NK cells being positively associated with bacterial burden drove PC1 (Figure 4D). One exception to these trends was found in a single male BCG vaccinated sample (Figure 4C).

We also noticed that Δ*lsr2* vaccinated mice lung samples were linearly separable across PC1 and PC2 (Figure 4B). We compared differences in protein expression between Δ*lsr2* vaccinated samples and all other samples to identify what cell-type-specific protein expression may be specific to Δ*lsr2* vaccination. Surprisingly, we found an overall increase in CD39 expression in both dendritic cells and granulocytes. This observed long term change in CD39 expression suggests that the high CD39 expression in dendritic cells from Δ*lsr2* inoculated mice described in our previous work may reflect a more sustained immune response to Δ*lsr2* inoculation compared with the other vaccine strategies (21).

### Unvaccinated mice show signs of CD4 T cell exhaustion 3 weeks post Mtb challenge

We next focused on cell-type-specific protein expression at ‘early infection’ (Figure 5A). As expected, pseudobulked PCA showed that PC1 (32.76% of variation in the dataset) is broadly separable by PBS control or vaccinated mice, with male PBS control mice distinctly grouped away from the other mice (Figure 5B). PC2 (21.09% of the variation in the dataset) separated PBS control and BCG vaccinated male and female mice from Δ*lsr2* vaccinated and *Mtb*+abx mice, except for a single male BCG mouse (Figure 5B). Thus, PC2 could indicate that the inoculation strain (*Mtb* vs. *M. bovis*) has a prolonged effect on immune reprogramming, after the similar strain effect observed at the ‘baseline’ timepoint.

**Figure 5.**
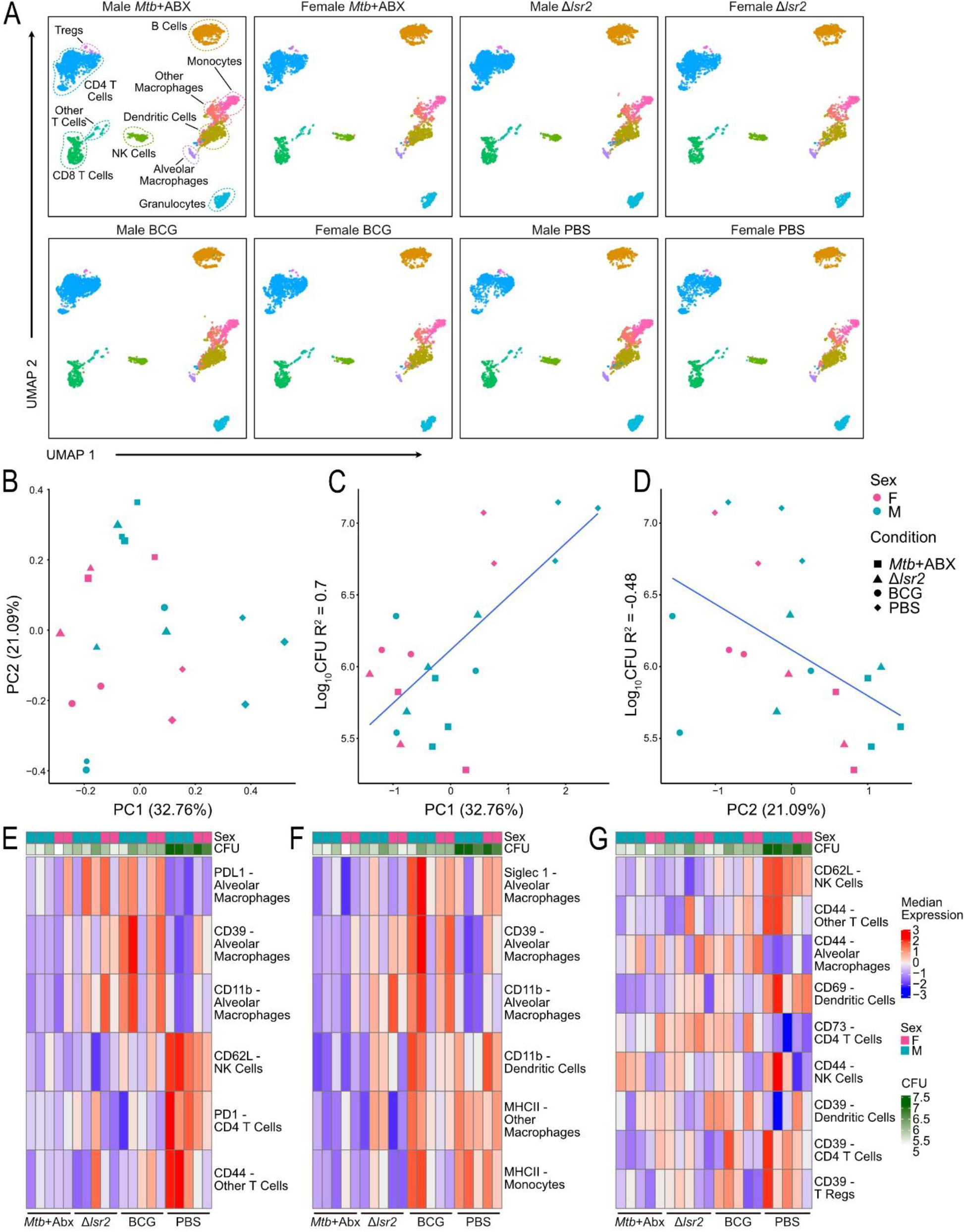
Immune profile of the male and female mouse lung 3 weeks after *Mtb* challenge of inoculated mice. (A) Uniform manifold approximation mapping (UMAP) showing the distribution of immune cells in the lungs based on their protein profiles, separated by treatment and sex. (B) Scatterplot of the first two principal components (PC) from a PC analysis of pseudobulked protein expression profiles for all samples. Shape represents condition and colour represents sex. (C-D) Scatterplot showing the relationship between (C) PC1 or (D) PC2 scores (x-axis) and bacterial burden (log_10_CFU, y-axis). Conditions are shape and sex is colour. (E-G) Heatmap of cell-type-specific protein expression driving (E) PC1, (F) PC2, and (G) proteins separating PBS males from other groups. Rows are cell types, columns are samples, and heat is the median expression of pseudobulked protein expression in each cell-type.

Next, we investigated cell-type-specific protein expression associated with PC1 (vaccinated vs. unvaccinated) and PC2 (BCG and PBS vs. Δ*lsr2* and *Mtb*+abx). PC1 scores were positively associated with bacterial burden, reflecting a higher overall *Mtb* CFU in PBS control samples (Figure 5C). PC1 scores were also different between conditions and sex. PC2 scores were potentially negatively associated with log_10_CFUs and associated with condition (Figure 5D).

Proteins driving PC1 expression during ‘early infection’ were CD62L in NK cells, PD1 expression in CD4 T cells, and CD44 expression in other T cells (Figure 5E). These cell-type-specific expression patterns were predominantly driven by male PBS control samples. We also found alveolar macrophage protein expression was associated with PC1, namely PDL1, CD39, and CD11b, which all had low expression in male PBS control samples and higher expression in BCG vaccinated samples (Figure 5E). Proteins driving PC2 were broadly driven by lower MHCII, CD39, and CD11b expression in macrophages, monocytes, and dendritic cells in the *Mtb*+abx and Δ*lsr2* groups compared to the BCG and PBS groups (Figure 5F).

When performing traditional differential protein expression analysis between PBS control male samples and all other samples (i.e., the samples driving the largest proportion of variance during ‘early infection’) we found increased activity of CD44 in NK cells and other T cells, CD39 in CD4 T cells and Tregs, CD62L in NK cells, and CD69 in dendritic cells (Figure 5G). We also found decreased activity of CD39 in dendritic cells, CD44 in macrophages, and CD73 in CD4 T cells in PBS control male mice during early *Mtb* challenge (Figure 5G).

### Immunological response during late Mtb infection was conditional on both sex and vaccine strategy, with male mice in the Mtb+abx group reflecting the unvaccinated condition

Lastly, we investigated cell-type-specific protein expression at ‘late infection’ (Figure 6A). Pseudobulked PCA showed that PC1 (45.09% of variation) is broadly separable by PBS control or vaccinated mice, as expected (Figure 6B). We also found that *Mtb*+abx male mice grouped more closely with PBS control mice (Figure 6B). This global cell-type-specific immune profile reflects the bacterial burden previously found in these mice, with *Mtb*+abx male mice having a comparable bacterial burden to PBS control male mice (Figure 1C). Interestingly, *Mtb*+abx female mice showed a global lung immune profile and lung bacterial burden in line with Δ*lsr2* and BCG vaccinated mice (Figure 6A-B, Figure 1C). PC1 (45.09% of the variation) is separated by vaccinated mice versus unvaccinated mice and male *Mtb*+abx. PC2 (22.73% of the variation) was linearly separable by sex, with PBS control males and females residing on the ends of PC2 (Figure 6B).

**Figure 6.**
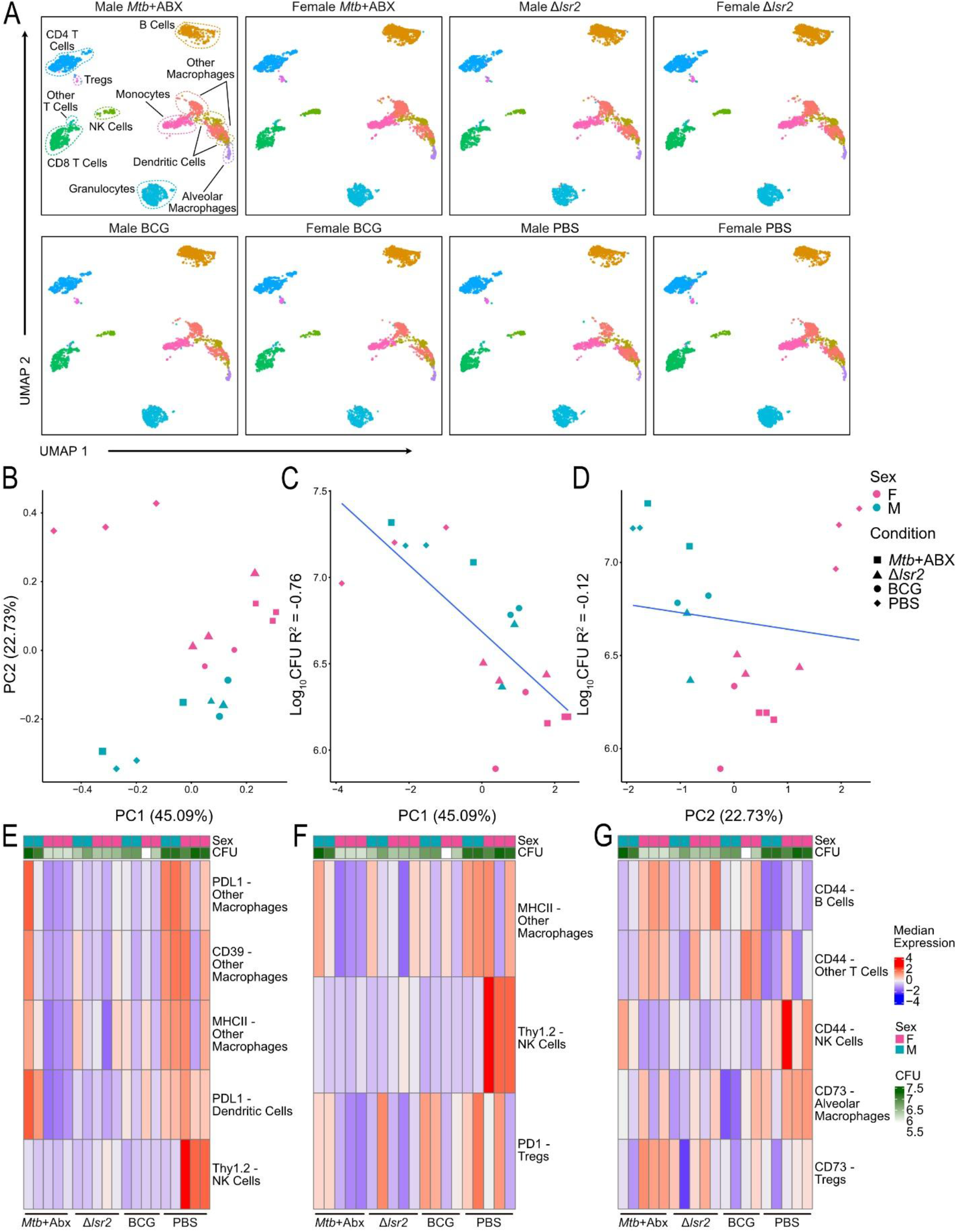
Immune profile of the male and female mouse lung 12 weeks after *Mtb* challenge of inoculated mice. (A) Uniform manifold approximation mapping (UMAP) showing the distribution of immune cells in the lungs based on their protein profiles, separated by treatment and sex. (B) Scatterplot of the first two principal components (PC) from a PC analysis of pseudobulked protein expression profiles for all samples. Shape represents treatment and colour represents sex. (C-D) Scatterplot showing the relationship between (C) PC1 or (D) PC2 scores (x-axis) and bacterial burden (log_10_CFU, y-axis). Conditions are shape and sex is colour. (E-G) Heatmap of cell-type-specific protein expression driving (E) PC1, (F) PC2, and (G) sex differences within inoculated conditions (i.e., *Mtb*+ABX, Δ*lsr2*, and BCG). Rows are cell types, columns are samples, and heat is the median expression of pseudobulked protein expression in each cell type.

We next sought to investigate cell-type-specific protein expression associated with PC1 (vaccinated vs. unvaccinated and male *Mtb*+abx). As expected, we found that PC1 negatively correlated with bacterial burden and was divergent between conditions (Figure 6C). Proteins driving PC1 at ‘late infection’ were Thy1.2 in NK cells (high in PBS control females), PDL1 in dendritic cells (high in PBS control samples and *Mtb*+abx males), and MHCII, PDL1, and CD39 in other macrophages (high in PBS control samples and *Mtb*+abx males; Figure 6E).

We then investigated PC2, which was sex-biased, associated with condition, and contained a condition-by-sex interaction driven by male *Mtb*+abx mice. PC2 does not correlate with bacterial burden because female PBS control mice are highly infected with *Mtb* (Figure 6D). Thy1.2 expression in NK cells and MHCII expression were associated with PC2 (Figure 6F). These proteins were also associated with PC1.

This pattern of global sex-biased cell-type-specific protein expression was not found at ‘baseline’ or ‘early infection’; thus, the male and female lung immune system likely undergoes different immune strategies from one another overtime. As such, while there is sex bias in *Mtb* response strategies, female mice only showed superior protection within the *Mtb*+abx (representing natural immunity) and BCG vaccinated groups (Figure 1F).

Female mice overall had a lower bacterial burden than males when considering all vaccine strategies together. Accordingly, we investigated sex differences in cell-type-specific protein expression among all vaccinated mice (Figure 6G). We found higher expression of CD44 in NK cells in vaccinated male mice and higher CD44 in NK cells in PBS control females. CD44 was higher in other T cells and B cells in vaccinated female mice. Lastly, CD73 may be higher in vaccinated female mice in both Tregs and macrophages. Accordingly, expression of these cell-type-specific proteins may contribute to the superior BCG vaccine-induced and natural protective immunity in female mice compared to males.

## Discussion

We performed the first global profiling of the mouse lung immune environment in response to various *Mtb* vaccination strategies. This design highlights the complexity of the host cellular immune ecosystem and the importance of including both male and female mice when defining immune factors that correlate with protection and prevention of TB. We used an analysis approach focusing on PCA rather than a series of pairwise cell-type-specific differential protein analyses. This PCA approach can have several key advantages when investigating datasets with many features (i.e., cell-type-specific proteins) and independent variables (i.e., vaccine strategy and sex). Firstly, the unsupervised nature of PCA allows for investigation of coordinated trends of cell-type-specific protein expression across all conditions rather than interpreting a list of individual pairwise comparisons (37,38). Secondly, the dimensionality reduction of PCA dramatically reduces the number of pairwise comparisons inherent to cell-type-specific differential expression, which can often mask key proteins driving the variance in the dataset (39). Thirdly, each PC provides the variance of the dataset explained (38). As such, PCA allowed us to measure the effect size of the cell-type-specific protein in relation to the variance in the entire dataset (e.g., these proteins drive 40% of the variability at the ‘baseline’ timepoint) (40), rather than providing that a protein’s median expression increases by 20%. As such, PCA-based approaches are extremely common in bulk and single-cell ‘omic analyses (38,40,41).

### The mouse lung immune system replicates the sex bias found in response to TB infections and vaccination prevalent in humans

In humans, males are disproportionately affected by TB and are 1.7 times more likely to develop active TB disease than females (12). This sex bias has also been found in mice, with male mice showing accelerated disease progression and increased mortality to *Mtb* infection (13). Recent studies have also shown a weaker or lack of protective efficacy from BCG in male mice compared to female mice (42,43). Our data recapitulates the sex bias in BCG vaccination response as well as the increased susceptibility of male mice to *Mtb* infection. In a recent study on BCG induced immunity, Parvathy et al. compared the sex bias of BCG and two recombinant BCG vaccines on their protective efficacy. They reported that BCG had the lowest protective efficacy in male mice and that the two recombinant vaccines only improved protection in males. However, all three vaccines performed equally in females and the recombinant vaccines had no effect on improving protection in female mice over BCG (43). Interestingly, we did not see this discrepancy in the Δ*lsr2* vaccinated group with both male and female mice having less variation in vaccine-induced immunity and a comparable level of protection to TB. While preliminary, we also propose one application of the Δ*lsr2* vaccine candidate could be to help decrease the sex-bias in the vaccination response.

### Unlike with BCG, Δlsr2 vaccination resulted in sustained, long-term changes in the lung immune system

Δ*lsr2*’s severe attenuation up to 18 weeks after vaccination provides further evidence that Δ*lsr2* is safe and a good live attenuated vaccine candidate (28). Δ*lsr2* had a low CFU of 100-1000 and was undetectable in 2/5 mice. With BCG, 4/5 mice had undetectable bacteria, and one mouse had a CFU <100 in the lungs after 18 weeks. One hypothesis for the lack of efficacy of BCG is that it is cleared from the host too quickly to induce long term protective immunity. A cytomegalovirus (CMV)-based vaccine provides proof of concept for the hypothesis that a long-acting vaccine could control *Mtb* infection (44). CMV establishes a persistent infection in humans, undergoing spontaneous reactivation cycles, and is unique in its ability to induce and maintain lifelong circulating tissue resident effector-differentiated CD4 and CD8 T cells (45). Rhesus macaques vaccinated with a rhesus CMV vector expressing 9 *Mtb* proteins reduced the overall extent of *Mtb* infection by 68% compared to unvaccinated macaques, with 41% showing no TB disease (44). Moreover, we detected immune system rewiring up to 18 weeks after Δ*lsr2* vaccination in mice, which was driven by CD44 in CD4 T cells and Tregs and CD39 in dendritic cells and granulocytes. It is important to note that there is a major concern of safety regarding CMV-based vaccines and the ability to attenuate the virus without impairing immunogenicity (45), thus making Δ*lsr2*’s severe attenuation, low sustained growth and long-term stimulation of the adaptive immune system an even more promising observation.

### Previously infected male mice show no natural immunity to TB by late-stage Mtb infection

In the *Mtb*+abx treatment group, we initially saw protection from natural immunity at the ‘early infection’ timepoint, however by ‘late infection’ there was none to little protective efficacy compared to unvaccinated PBS controls. This data recapitulates the findings from the study by Henao-Tamayo et al., which used a similar methodology (32). Epidemiological data in TB endemic regions also indicates that people who had TB once are at four times higher risk of developing TB when reinfected than those who are primarily infected (46). Moreover, we saw the greatest sex-bias in the *Mtb*+abx treatment group at the ‘late infection’ timepoint, with males having a higher bacterial burden than female mice. The immune environment in these severely infected *Mtb*+abx male mice was defined by a low abundance of B cells, NK cells, Tregs and other T cells, and a high abundance of granulocytes and CD8 T cells. This is consistent with previous studies that have reported male mice to display an exaggerated inflammatory response to *Mtb* and smaller B cell follicles in the lungs, which both correspond to increased susceptibility and mortality (13,47). However, unlike Dibbern et al. that reported a lack of neutrophil accumulation in the lungs as part of the increased inflammatory response, we observed an increased abundance of granulocytes in *Mtb*+abx males.

### CD39 correlates with infection severity in a cell-type and infection-stage dependent manner

CD4 T cell infiltration in the lungs was associated with protection post *Mtb* challenge. However, PD1 expression on CD4 T cells was positively associated with unvaccinated mice and increased bacterial burden during ‘early infection’. Previous work has shown that the PD1-PDL1 pathway impairs protective T cell function during late stage BCG infection and enables bacterial persistence (48). PD1 has also been associated with CD39 expression as markers of T cell exhaustion in cancer models (49), and we previously reported on the correlation between high CD39 expression and higher *Mtb* bacterial burden (21). Unlike the cancer models, in our previous work we did not find a pattern of PD1 and CD39 co-expression on T cells correlating to high *Mtb* bacterial burden (21). However, here we did find that both PDL1 and CD39 are strongly associated with bacterial burden during ‘late infection’ in dendritic cells and macrophages. Taken together, this data highlights the need for non-exhausted T cells for bacterial control and further supports the importance of defining the functional state of induced T cells to determine their correlation to protection.

### Thy1.2 expression on NK cells drives improved protection of female mice during late-stage infection

Interestingly, NK cells consistently showed up as correlated with lower *Mtb* bacterial burden, particularly at the ‘late infection’ timepoint and in female mice. This is consistent with previous work linking NK cell expansion to protection after mucosal and intravenous BCG vaccination (50,51), as well as another study linking high NK cell abundance to low lung pathology and controlled TB disease in non-human primates (52). Unlike in previous work, we saw a strong female sex-bias in the NK cell response. Even with unvaccinated mice at ‘late infection’, where male and female mice had comparable CFUs, female mice showed a much higher Thy1.2 expression on NK cells reflecting a substantial increase in NK cell activation (53,54). To our knowledge, only one recent study has previously seen a sex-bias in the NK cell response to *Mtb* infection. Hertz et al. used RAG2 knockout mice to investigate sex differences in the innate immune response to *Mtb* and found that in the absence of T and B cells, the expected response to an *Mtb* infection by male and female mice was flipped. Female RAG2 knockout mice exhibiting an elevated inflammatory response and were more susceptible to *Mtb* (14). Male mice, on the other hand, showed greater protection to *Mtb* which was correlated with a more robust NK cell response (14). Taken together, this work underscores the need for a better understanding of the differing interplay between innate and adaptive TB immunity between sexes. We further showed that by ‘late infection’ male and female mice showed highly different immune strategies in response to *Mtb* infection. This pattern included the differing NK cell response as well as CD44 expression on diverse immune cell-types including B cells, and other T cells. Our work builds on the growing appreciation for the complexity of *Mtb*- and TB vaccine-mediated immunity as well as the broader differences in TB disease progression between males and females.

## Limitations and future directions

The complexity inherent to *Mtb* infection and vaccination warranted a broad, systematic view thus we chose to include a breadth of conditions, timepoints, and sexes in one CyTOF *Mtb* challenge experiment. CyTOF allows for 20 samples to be multiplexed within a single experimental run. While this allows us to investigate multiple conditions and sexes without counting for experimental batch effects, this did limit the sample size for certain variables such as sex. However, the breadth of conditions within a single experimental batch we were able to capture enables future data reanalysis depending on the biological question. Moreover, experiments of this design are exploratory in nature, meaning that many individual findings (e.g., Δ*lsr2* vaccinated mice showing similar improved protection across sexes) are subject to replication and validation. Our replication of many previous findings and well established *Mtb* immunity (e.g. the correlation of CD4 T cells with reduced bacterial burden) serves to validate our study design and contextualize this work within a single system to provide a foundation for developing hypotheses for downstream interrogation.

CyTOF provides the distinct advantage of multiplexing samples, studying protein expression, and having a high number of cells per sample. However, it does not allow for the discovery of new markers or pathways in the way that methods such as scRNA-seq have. As such, follow-up work using single-cell and spatial transcriptomic approaches may help unveil which specific biological systems and pathways are driving the patterns we found in this study. Unlike CyTOF which relies on species-specific antibodies, single-cell genomic experiments are also species agnostic, allowing a broader range of animals to be investigated.

This study provides proof of concept that our single-cell proteomics design can model the complex ecosystem of *Mtb*- and vaccine-induced immunity with multiple confounders including sex, vaccination status, and prior exposure/treatment, and capture variance and robust trends not previously reported in *Mtb* infection. We highlighted the continued presence of immune system rewiring after long-term vaccination, the importance of non-exhausted T cells during ‘early infection’ for bacterial control, and the presence of sex- and vaccine-strategy specific immunological strategies in response to *Mtb* challenge. This work strongly encourages more emphasis be placed on understanding sex-based immune response differences for the development of successful next generation TB vaccines and treatments. This work also provides an unbiased database that allows researchers to interrogate patterns of cell-type abundance and cell-type protein expression across various *Mtb* infection outcomes.

## Acknowledgements

Thank you to the wonderful staff at the Division of Comparative Medicine and the Toronto High Containment Facility. We also wish to acknowledge Dr. Blair Gordon for generating the *Mtb* Δ*lsr2* mutant, Dr. Jesse Gillis for his critical advice on ‘omics statistics and incorporating MetaNeighbor analysis, and Dr. Wenxi Xu for his advice on cluster cell-type naming. The study was supported by Canadian Institutes for Health Research Funds PJT-156261 and PJT-186285 (J.L.). D.J.S. is supported by the NSERC-PDF scholarship and the McLaughlin Centre Scholars Grant. Animal figures were created with BioRender.com.

## Author contributions

Conceptualization J.W., D.S. and J.L.; Methodology J.W. and D.S.; Investigation J.W and D.S.; Visualization J.W. and D.S.; Funding acquisition J.L.; Project administration J.W.; Supervision J.L.; Writing – original draft J.W., D.S. and J.L.; Writing – review and editing J.W. and J.L.

## Declaration of interests

“Authors declare that they have no competing interests.”

## Methods

| Antibodies | Source | Identifier |
| --- | --- | --- |
| **Antibodies** |  |  |
| 141Pr anti-mouse Ly-6G/C/Gr-1 (clone RB6-8C5) | Fluidigm | Maxpar Mouse Spleen/Lymph Node Phenotyping Panel Kit, Cat#201306; RRID: not available |
| 142Nd anti-mouse CD39 (clone 24DMS1) | Fluidigm | Cat#3142005C; RRID: not available |
| 144Nd anti-mouse MHC Class I (clone 28-14-8) | Fluidigm | Cat#3144016C; RRID: not available |
| 145Nd anti-mouse CD69 (clone H1.2F3) | Fluidigm | Maxpar Mouse Spleen/Lymph Node Phenotyping Panel Kit, Cat#201306; RRID: not available |
| 146Nd anti-mouse F4/80 (clone BM8) | Fluidigm | Cat#3146008C; RRID: not available |
| 147Sm anti-mouse CD45 (clone 30-F11) | Fluidigm | Maxpar Mouse Spleen/Lymph Node Phenotyping Panel Kit, Cat#201306; RRID: not available |
| 148Nd anti-mouse CD11b/Mac-1 (clone M1/70) | Fluidigm | Maxpar Mouse Spleen/Lymph Node Phenotyping Panel Kit, Cat#201306; RRID: not available |
| 149Sm anti-mouse CD19 (clone 6D5) | Fluidigm | Maxpar Mouse Spleen/Lymph Node Phenotyping Panel Kit, Cat#201306; RRID: not available |
| 150Nd anti-mouse Ly-6C (clone HK1.4) | Fluidigm | Cat#3150010C; RRID: not available |
| 151Eu anti-mouse CD25 (clone 3C7) | Fluidigm | Maxpar Mouse Spleen/Lymph Node Phenotyping Panel Kit, Cat#201306; RRID: not available |
| 152Sm anti-mouse CD3e (clone 145-2C11) | Fluidigm | Maxpar Mouse Spleen/Lymph Node Phenotyping Panel Kit, Cat#201306; RRID: not available |
| 153Eu anti-mouse CD274/PD-L1 (clone MIH5) | Fluidigm | Cat#3153031C; RRID: not available |
| 154Sm anti-mouse CD73 (clone TY/11.8) | Fluidigm | Cat#3154019C; RRID: not available |
| 156Gd anti-mouse CD90.2/Thy-1.2 (clone 30-H12) | Fluidigm | Cat#3156006C; RRID: not available |
| 158Gd anti-mouse Foxp3 (clone FJK-16s) | Fluidigm | Cat#3158003C; RRID: not available |
| 159Tb anti-mouse CD279 (clone RMP1-30) | Fluidigm | Cat#3159006C; RRID: not available |
| 160Gd anti-mouse CD62L (clone MEL-14) | Fluidigm | Maxpar Mouse Spleen/Lymph Node Phenotyping Panel Kit, Cat#201306; RRID: not available |
| 161Dy anti-Ki-67 (clone B56) | Fluidigm | Cat#3161007C; RRID: not available |
| 162Dy anti-mouse CD366/TIM-3 (clone RMT3-23) | Fluidigm | Cat#3162029C; RRID: not available |
| 168Er anti-mouse CD8a (clone 53-6.7) | Fluidigm | Maxpar Mouse Spleen/Lymph Node Phenotyping Panel Kit, Cat#201306; RRID: not available |
| 169Tm anti-mouse TCRβ (clone H57-597) | Fluidigm | Maxpar Mouse Spleen/Lymph Node Phenotyping Panel Kit, Cat#201306; RRID: not available |
| 170Er anti-mouse CD169/Siglec-1 (clone 3D6.112) | Fluidigm | Cat#3170018C; RRID: not available |
| 171Yb anti-mouse CD44 (clone IM7) | Fluidigm | Maxpar Mouse Spleen/Lymph Node Phenotyping Panel Kit, Cat#201306; RRID: not available |
| 172Yb anti-mouse CD4 (clone RM4-5) | Fluidigm | Maxpar Mouse Spleen/Lymph Node Phenotyping Panel Kit, Cat#201306; RRID: not available |
| 173Yb anti-human/mouse Granzyme B (clone GB11) | Fluidigm | Cat#3173006C; RRID: not available |
| 174Yb anti-mouse I-A/I-E (clone M5/114.15.2) | Fluidigm | Cat#3174003C; RRID: not available |
| 176Yb anti-mouse B220 (clone RA3-6B2) | Fluidigm | Maxpar Mouse Spleen/Lymph Node Phenotyping Panel Kit, Cat#201306; RRID: not available |
| 209Bi anti-mouse CD11c (clone N418) | Fluidigm | Cat#3209005C; RRID: not available |

### Animals

Male and female C57BL/6 mice between 6-10 weeks of age were purchased from Charles River Laboratories. Experiments involving *Mycobacterium tuberculosis* were conducted in the high containment facility, biosafety level III laboratory at the University of Toronto. Animal studies and procedures were approved by the University of Toronto Animal Care Committee and performed in accordance with the committees’ ethical standards.

### Bacterial strains and culture conditions

*Mycobacterium tuberculosis* H37Rv, *Mtb* Δ*lsr2*, and *M. bovis* BCG-Japan were prepared and titrated as previously described (55). Briefly, *Mtb* strains were grown at 37°C in Middlebrook 7H9 broth (BD) supplemented with 0.2% glycerol, 10% albumin-dextrose-catalase, and 0.05% Tween 80 or on 7H10 agar supplemented with 1% casamino acids (Life), 0.5% glycerol, and 10% albumin-dextrose-catalase. Hygromycin (BioShop) was added at a concentration of 75ug/mL for *Mtb* Δ*lsr2*. BCG was grown on 7H10 agar supplemented with 1% casamino acids (Life), 0.5% glycerol, and 10% oleic acid-albumin-dextrose-catalase or at 37°C in Middlebrook 7H9 broth (BD) supplemented with 0.2% glycerol, 10% albumin-dextrose-catalase, and 0.05% Tween 80.

### C57BL/6 mice infection with mycobacterium

For aerosol infections, ∼100 colony-forming units (CFUs) of *Mtb* H37Rv per mouse was delivered using a Glas-Col nebulizer Inhalation Exposure System (55). Delivery dosage was confirmed by quantifying CFU in the lungs 1 day after exposure in 4 control mice. At various timepoints after inoculation (18 weeks post-vaccination, pre-challenge with *Mtb* – ‘baseline’; 3 weeks post-challenge with *Mtb* – ‘early infection; 12 weeks post-challenge with *Mtb* – ‘late infection’) lungs and spleen CFUs were determined. Lungs and spleen were homogenized in PBS and tissue homogenate was serial diluted and coated on 7H10 agar plates. CFUs were counted on the plates after a 4-week incubation at 37°C. Tissues were also collected for CyTOF analysis or fixed in 10% neutral buffered formalin (Sigma Aldrich) for histological analysis.

### Intranasal vaccination of C57BL/6 mice

5×10^3^ CFU *Mtb* H37Rv, 5×10^5^ CFU *Mtb* Δ*lsr2*, or 5×10^5^ CFU *M. bovis* BCG-Japan were delivered intranasally in 20μL PBS. 18 weeks after vaccination, mice were aerosol infected with *Mtb* as described above.

### Antibiotic treatment of *Mtb* infected C57BL/6 mice

4 weeks after initial intranasal inoculation with *Mtb* H37Rv, as described above, mice were treated with rifampicin (100mg/L) and isoniazid (200mg/L) in drinking water for 12 weeks. Mice were then rested for 2 weeks to allow residual drug to wash out before they were aerosol challenged with *Mtb* as described above.

### Tissue isolation

Lung and spleen tissue was isolated as previously described (56). Lungs and spleen were minced into fine pieces then digested in RPMI 1640 medium without calcium/magnesium (Gibco) containing 10% FBS, 1% HEPES (Life), 1 mg/mL Collagenase from Clostridium histolyticum (Sigma) and 0.15 mg/mL DNase I (Sigma) on a shaker at 37°C for 1 hour with the lungs or 30 min with the spleen. Digested tissue was filtered through a 70 μm cell strainer to obtain single-cell suspensions and an aliquot was plated on 7H10 agar plates. CFUs were counted after a 4-week incubation at 37°C. The remaining lung and spleen single cell suspensions were red cell-lysed and then processed for CyTOF analysis.

### Time-of-Flight mass cytometry (CyTOF)

Details for each antibody used for CyTOF are listed in Table 4.1. Single cell suspensions from individual samples were washed with PBS and stained with 1μM Cisplatin in PBS for 5min then quenched with Maxpar® Cell Staining Buffer and FBS underlay. Samples were barcoded using Cell-ID™ 20-Plex Pd Barcoding Kit (Standard Biotools) according to manufacturer’s instructions. Barcoded samples were then combined and resuspended in Maxpar® Cell Staining Buffer containing metal-tagged surface antibodies and Fc block (CD16/32; Bioxcell) for 30 minutes at 4°C. Cells were then washed, fixed, and permeabilized (Intracellular cytokine buffer kit and Foxp3 staining kit, eBioscience) before being stained with metal tagged intracellular antibodies for 30 minutes at room temperature. Directly conjugated antibodies were purchased from Standard Biotools. Finally, cells were incubated overnight in PBS containing 0.3% (w/v) saponin, 1.6% (v/v) paraformaldehyde (Polysciences Inc) and 50nM Iridium (Standard Biotools). Cells were analyzed on a Helios2 mass cytometer (Standard Biotools) at The Hospital for Sick Children Center for Advanced Single Cell Analysis.

### CyTOF data processing

In this study, three CyTOF datasets were generated that explore various aspects of the immunology dynamics following an *Mtb* challenge after various vaccine strategies in male and female mice. To note, one female PBS lung sample, an uninfected control, was excluded from the ‘baseline’ (18 weeks post-vaccination, pre-challenge with *Mtb*) CyTOF analysis as the staining did not work. CyTOF data processing was applied to each timepoint individually and followed the same workflow and pipeline as previously described (21). Briefly, output fcs files were debarcoded using the “CATALYST” R package (57,58) as well as the debarcoding template “debarcoding_gene_template.csv” (Standard Biotools Cell-ID™ 20-Plex Pd Barcoding Kit). The prepData() function in CATALYST was then used with default parameters to filter non-mass channels and perform an arcsinh-transformation (cofactor = 5) (57,58). A SingeCellExperiment object, an S4 object built to be compatible with many single-cell ‘omics R packages, was generated (59). Then, assignPrelim() was used to assign each cell to its sample of origin before filtering cells that were not assigned a unique barcode (57,58). The estCutoffs(), plotYields(), and applyCutoffs() functions were used to estimate barcode distributions, plot distributions, and filter cells whose distributions suggest technical artifacts such as doublets, respectively (57,58). Normalized and debarcoded sample files were saved as their own .fcs files.

For each dataset, data normalization, visualization, and quality control was performed by adapting the “cytofWorkflow” R package and module to our experimental design (60). This workflow predominantly relies on the “flowCore”, “CATALYST”, and “diffCyt” R packages (57,58,61). The following analysis was applied to each dataset individually. Debarcoded .fcs files for each sample were loaded into R assigned the mass channel and mass key between each antibody to the biological symbol of the antibody (e.g., Tag 142Nd = CD39) (62). Quality control of these combined and normalized data was done with the plotExprs() function, plotCounts(), and pbMDS() functions (62). The plotNRS() function was used to visualize non-redundancy scores, which measure protein variance between cell types and conditions (62).

Normalized and quality-controlled data were then clustered to generate cell types. All proteins were used to cluster the cells, rather than exclusively proteins known to distinguish cell types because including cell-state proteins allows us to empirically generate cellular subclusters that are driven by cellular activation, exhaustion, or apoptosis (57,58). Cells were clustered in each dataset using the cluster() function in the “CATALYST” R package, which first generates a 10 by 10 grid of clusters using FlowSOM (63), defined as self-organized map clustering optimized for CyTOF data. The cluster() function then groups these 100 clusters into 30 metaclusters using the ConsensusClusterPlus() R package (64). A heatmap that plots protein expression to each cluster using the “plotExprHeatmap” was generated and used to manually assign cell-types to each cluster. Visualization of the relationship between protein expression, cluster ID, and sample-level cell-type proportions was done with the plotMultiHeatmap”. Dimed tSNE and UMAP (65) dimensionality reductions were performed with the runDR() functions. tSNE and UMAPs showing cell-type, cluster type, and protein expression are all plotted with the plotDR() function; for computational efficiency, 1800 randomly selected cells from each sample were used to generate the tSNE and UMAPs. In each dataset, cells characterized by non-specific protein binding (undefined) or with contradicting cell-type proteins (doublets) were removed from downstream analysis.

Cluster/cell-type proportions were calculated using the diffcyt() R package with the “DA” module (61). Pseudobulked cluster/cell-type-specific gene expression using the diffcyt() R package with the “DE” module were also generated (61). The final SingleCellExperiment, cell-type expression matrix and cell-type proportion matrix for each dataset were saved for dataset-specific downstream analysis.

### Cell-type protein mapping between CyTOF timepoints

The “MetaNeighbor” R package (66) was used to match cell-types (i.e., ‘baseline’ vs. ‘early infection’, ‘baseline’ vs. ‘late infection’, ‘early infection’ vs. ‘late infection’). The MetaNeighborUS() function was used to calculate a cell-by-cell area under the operating curve (AUROC) matrix between each timepoint (66). The AUROC between row ‘i’ and column ‘j’ represents the likelihood that cells mapping to that cell-type in each timepoint are from the same population. This AUROC matrix was plotted using the heatmap.2 function to visualize the distribution of cell identity similarity between timepoints. The topHits() function was then used to extract the cell-type pairs with an AUROC > 0.85. Lastly, matching pairwise cell-types (i.e., AUROC > 0.85) were collapsed into a graph, where nodes represented the cell-type at each timepoint and the edge represented the AUROC between each timepoint, to identify cell-types that matched between all three timepoints (66). This analysis was repeated at the metacluster and cluster levels.

### Differential cell-type abundance analysis with CyTOF data

Cell-type proportions were globally visualized between all three timepoints using the “ComplexHeatmap” R package (DOI: 10.18129/B9.bioc.ComplexHeatmap). The Heatmap() function was used to plot each cell-type in each sample, with samples annotated by log_10_CFU, sex, condition, and timepoint. The relationship between cell-type composition, sex, condition, and timepoint on log_10_CFU was compared with a factorial ANOVA (log_10_CFU ∼ cell-type composition + timepoint + Sex + condition). The relationship between cell-type composition and timepoint, sex, and condition was compared using a multi-way ANOVA (cell-type ∼ timepoint + sex + condition + timepoint*sex + timepoint*condition + sex*condition).

PCs that were associated with log_10_CFUs (FDR-adjusted p-value < 0.05), significantly different between conditions, and/or showed a sex-by-condition interaction were selected to be associated with *Mtb* vaccination. These PCs were then plotted against the log_10_CFU in each mouse, visualizing sex and condition, using the ggplot() function in the “ggplot2” R package function. The PC loadings for vaccination-associated PCs were then extracted and the cell-type-specific protein expression driving that PC were identified (|loading| > mean(|loading|) + sd(|loading|)) (67).

Within each timepoint, cell-type proportion was associated with log_10_CFUs using Spearman’s correlation. Cell-type proportion was also compared between sex and condition using a two-way ANOVA (proportion ∼ condition + sex + condition*sex). Like the PCA, cell-types that were associated with log_10_CFUs (FDR-adjusted p-value < 0.05), significantly different between conditions, and/or showed a sex-by-condition interaction, were selected to be associated with *Mtb* vaccination. This analysis was performed for cell-types, metaclusters, and clusters.

## Quantification and statistical analysis

Graphs and statistics were generated with GraphPad Prism v10 (GraphPad Software). Number of animals per group are described in the figure legends. Error bars represent the standard error of the mean (SEM). Data shown in figures with log-scale y-axis was transformed (log_10_) to conform to normality. Student’s t-test, one-way and two-way ANOVA with Sidak’s or Tukey multiple comparison test were used to determine statistical significance where appropriate. Significance is indicated by: * = p < 0.05, ** = p < 0.01, *** = p < 0.001, **** = p < 0.0001. ns = not significant.

## Notes

### Competing Interest Statement

The authors have declared no competing interest.

