## Supplemental Figure for "In-Depth Immune Profiling Reveals Cell-Type, Sex and Infection-Stage Dependent Differences for Tuberculosis Control"

### Supplemental figures

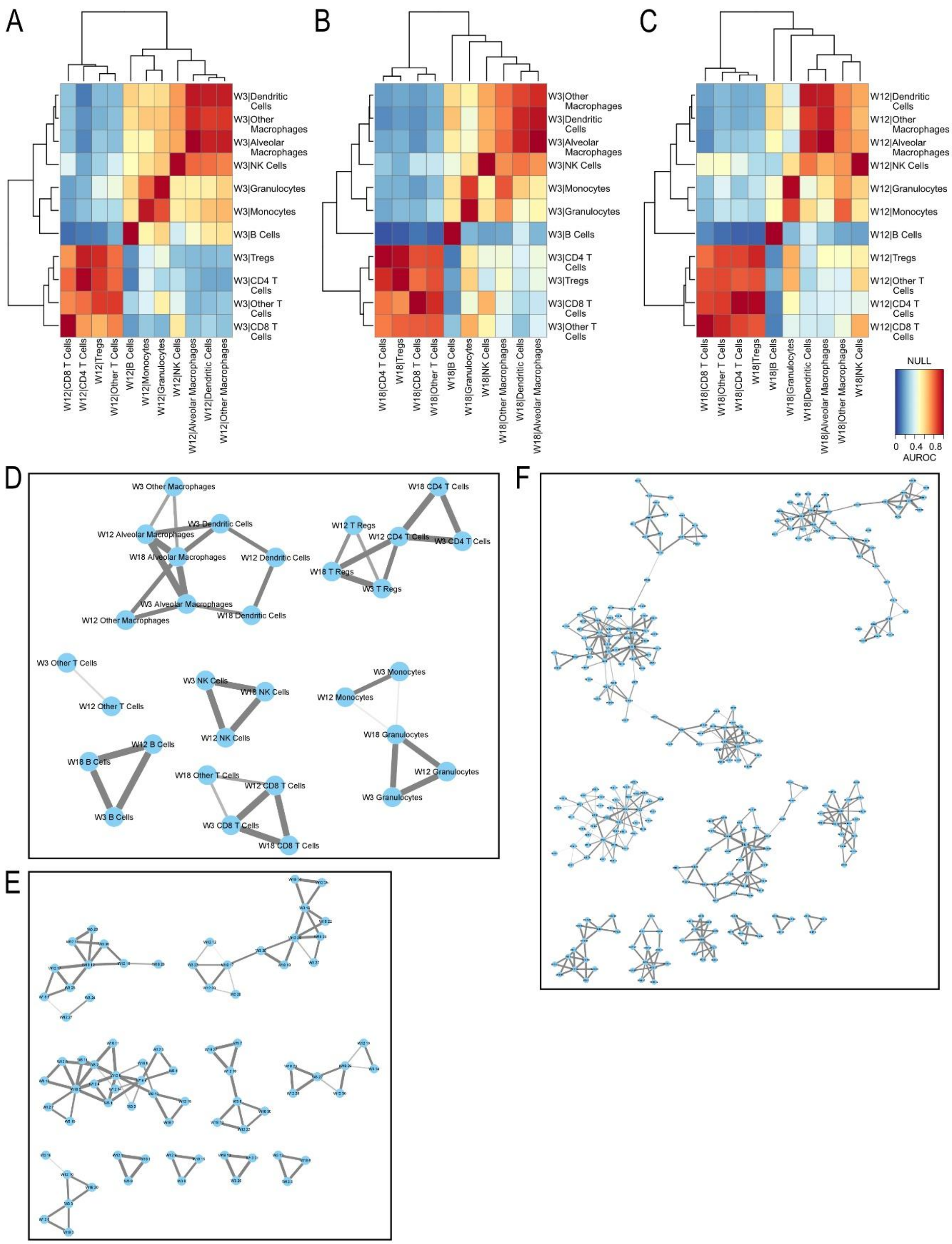


**Supplementary Figure S1 Cross-timepoint cell-cell relationships identified by MetaNeighbor.** (A) Heatmap representing cell-cell relationships between 'early infection' (W3) and 'late infection' (W12). (B) Heatmap representing cell-cell relationships between 'early infection' (W3) and 'baseline' (W18). (C) Heatmap representing cell-cell relationships between 'late infection' (W12) and 'baseline' (W18). Rows are cell-types called in the first timepoint listed, columns are cell-types called in the second timepoint listed, and heat is populated by the AUROC score mating each cell-type. (D-F) Graphical relationship between cell-types at 'early infection' (W3), 'late infection' (W12), and 'baseline' (W18). Each node is a (D) cell-type, (E) metacluster, and (F) cluster at one of the three timepoints. Edges represent a reciprocal top hit or an AUROC > 0.9 between nodes. A network of three nodes, where each node is one timepoint, represents a perfect one-to-one-to-one corresponding cell-type (e.g., B cells, NK cells). 'Early infection' is 3W post *Mtb* challenge; 'late infection' is 12W post *Mtb* challenge; and 'baseline' is 18W post-vaccination.
